# Scale Invariance in fNIRS as a Measurement of Cognitive Load

**DOI:** 10.1101/2021.08.31.458427

**Authors:** Chu Zhuang, Kimberly L. Meidenbauer, Omid Kardan, Andrew J. Stier, Kyoung Whan Choe, Carlos Cardenas-Iniguez, Theodore J. Huppert, Marc G. Berman

## Abstract

Scale invariant neural dynamics are a relatively new but effective means of measuring changes in brain states as a result of varied cognitive load and task difficulty. This study is the first to test whether scale invariance (as measured by the Hurst exponent, *H*) can be used with functional near-infrared spectroscopy (fNIRS) to quantify cognitive load. We analyzed *H* extracted from the fNIRS time series while participants completed an N-back working memory task. Consistent with what has been demonstrated in fMRI, the current results showed that scale-invariance analysis significantly differentiated between task and rest periods as calculated from both oxy- (HbO) and deoxy-hemoglobin (HbR) concentration changes. Results from both channel-averaged *H* and a multivariate partial least squares approach (Task PLS) demonstrated higher *H* during the 1-back task than the 2-back task. These results were stronger for *H* derived from HbR than from HbO. As fNIRS is relatively portable and robust to motion-related artifacts, these preliminary results shed light on the promising future of measuring cognitive load in real life settings.

**Author Summary:** Scale invariance reflects a pattern of self-similarity (or fractalness) across a time series of brain data. In human neuroscience studies using EEG and fMRI, higher scale invariance has been associated with individuals being in a state of minimal cognitive effort or while performing a relatively easy task compared to doing something more challenging. Functional near-infrared spectroscopy (fNIRS) is a flexible neuroimaging technique that can be used in naturalistic settings and measures the same underlying biological signal as fMRI. We expected that, if scale invariant brain states are indeed robust indicators of cognitive load or task difficulty, we should be able to replicate previous findings in fNIRS. Consistent with this hypothesis, we find that more scale invariant brain states are indeed associated with less cognitively demanding and more restful brain states in fNIRS data. This finding opens up a wide array of potential applications for monitoring cognitive load and fatigue in real-life settings, such as during driving, learning in schools, or during interpersonal interactions.

## 1. Introduction

Functional near-infrared spectroscopy (fNIRS) is a neuroimaging technique that has gained increasing attention in recent years due to relative robustness to motion artifacts and environmental noise, making it more suitable for neuroimaging outside standard laboratory settings [1,2] and with hard-to-test populations [3]. Like functional MRI, fNIRS measures changes in the brain’s hemodynamic response, but does so using light spectroscopy at near-infrared wavelengths. At the onset of regional neural activity, metabolic demands rise, and blood flow increases in that area. This increased blood flow leads to higher concentrations of oxygenated hemoglobin concentrations (HbO) and lower concentrations of deoxyhemoglobin concentrations (HbR). Measuring these oxy- and deoxy-hemoglobin concentration changes in fNIRS provide a complementary method to fMRI with lower cost and greater use-case flexibility [4]. Many findings from fMRI have been replicated using fNIRS, such as activation changes that result from varying cognitive load in working memory tasks [5–7].

Cognitive load refers to the level of demand and difficulty people bear when performing a cognitive task [8,9]. More specifically, it often indicates the amount of working memory used or the number of items people are actively holding in working memory [6,7]. However, when people are continuously under high cognitive load and there are sustained demands on attention and working memory, performance can decrease, and people may suffer from high levels of mental fatigue. This fatigue can be very dangerous, especially for those who work in positions requiring sustained attention, such as pilots, doctors, technicians under military duties etc [10,11]. Similarly, in an educational setting, children may not learn effectively or perform well if they are already experiencing high levels of cognitive fatigue[12].

Both behavioral (i.e., speed and accuracy in a task) and physiological measures (i.e., skin conductance, blood glucose, cardiac function) have been used to measure cognitive load in previous work [13,14]. However, behavioral measures do not directly reflect cognitive load and may neglect ‘compensatory effort’[15]. That is, people may exert extra effort and experience higher cognitive load while maintaining the same level of performance when they are under fatigue and distraction. Physiological methods are not able to index cognitive load separately from other processes, as they also measure cardiovascular and sympathetic nervous system responses evoked by processes unrelated to cognitive load, such as arousal [16] and stress [17,18]. In particular, when a person is experiencing mental fatigue, it can be hard to tell what changes reflect cognitive effort and what are due to arousal and stress related to performance, for example. Thus, physiological methods are neither specific nor fully sensitive to changes in cognitive load. To better monitor levels of cognitive load in these real-life situations and tailor interventions to mitigate fatigue, it would be advantageous to use mobile neuroimaging technology with a reliable neural measure of cognitive difficulty, effort, or fatigue.

Previous neuroimaging studies investigating cognitive load with fNIRS and fMRI have often centered around localized activation that differs based on task difficulty or working memory load [19–21]. However, the present study aimed to measure levels of cognitive load through a whole brain neural signature, *scale invariance* of the broadband brain signal, quantified by the Hurst exponent (*H*). This signature has been validated with fMRI and EEG studies and has proved sensitive and specific to changes of cognitive load [15,22,23]. Compared with fMRI and EEG, fNIRS is more robust to motion artifacts and environmental noise and is more portable, making it ideal for investigations in ecologically valid settings (for example, measuring fatigue during driving [24], or learning in school aged children [25].

Scale invariance (also called scale-free or fractal scaling) refers to a property of a signal wherein the same pattern appears across different scales. Specifically, in the temporal domain, scale invariance means that a time series signal has a persistent autocorrelation with long-memory dependence across timescales. Information of the temporal signal is contained in all frequencies and timescales, and no specific frequency band plays a dominant role. Therefore, from the frequency perspective, scale invariant neural signals exhibit a power-law relationship between Power Spectral Density of the signal (PSD), and its frequency (f), PSD(f)∼f-β, β>=0. When the signal is scale-invariant, the slope of this function, -β, should be negative (PSD close to 1/f), and hence β should be positive. The more scale-invariant the signal is, the more inclined the slope is and the higher value of β [22].

According to this formula, the degree of temporal scale invariance can be calculated by the Hurst exponent (*H*), which is related to β via the equation H=(β+1)/2, or β=2H-1. Therefore, if a signal is scale invariant, β should be positive and *H* should be larger than 0.5; and a higher value of *H*, corresponding with a higher value of β, indicates more scale-invariant signal. Previous research has found that *H*, the scale invariance index, is a robust neural indicator of cognitive load [22,26,27]. The effectiveness of *H* in quantifying cognitive load and task engagement has been validated in both fMRI (BOLD signals) and EEG studies (oscillatory activity, [15]. *H* decreases globally with changes in cognitive load and task difficulty, where it is highest during rest, lower while participants perform tasks, and shows the most suppression during highly cognitively demanding tasks. However, while the effectiveness of the Hurst exponent as a cognitive load measurement has been validated in both fMRI and EEG studies, it has not yet been examined in fNIRS. The present study addresses whether this temporal neural signature is also feasible in quantifying cognitive load with fNIRS.

### Study Design and Hypothesis

Based on previous findings in fMRI and EEG studies [22,26–28], we hypothesize that increasing levels of cognitive load will be associated with suppression of *H*, indicating decreased scale invariance as measured by fNIRS signals. To test this in the current study, we examined *H* in an N-back working memory task while fNIRS data were recorded.

The N-back task is a classic working memory paradigm which places high demands on working memory and attention with varying levels of task difficulty and cognitive load [6,7]. In the N-back working memory task, participants are required to match the current word/stimulus with the word presented in the previous Nth trial and make a response. The task’s difficulty is varied by the different Nth preceding trial participants required to remember and compare with (N=1,2,3), defined as ‘1-back’, ‘2-back’ and ‘3-back’ conditions. Here, the ‘1-back’ is the easiest and the ‘3-back’ is the most difficult. Accordingly, we hypothesize that scale invariance of fNIRS signal, measured as Hurst exponent (*H*), will decline with the increasing task difficulty: whereby the ‘1-back’ will show the highest *H*, while the ‘3-back’ will have the most suppressed/lowest *H*, indicating a higher cognitive load.

Results comparing task and rest conditions show that, as expected, *H* values averaged across all channels were higher during rest than during task, and higher in the 1-back task than the 2-back task. Contrary to our hypotheses, there were no significant differences between 3-back and other N-back levels, though this is consistent with the task-activation results examined in this same dataset reported in Meidenbauer et al. [29] and are likely due to participants disengaging with the task as indicated by poor task performance on the 3-back. These effects were somewhat larger for *H* calculated from the deoxyhemoglobin signal (HbR) than the oxyhemoglobin signal (HbO), which may be due to the tighter coupling between HbR and the BOLD response observed in fMRI relative to HbO [1]. In addition, a multivariate partial least squares analysis (PLS) which examined N-back level differences in *H* values in each channel replicated this effect, where higher *H* values were found for the 1-back versus the 2-back task, and the effect was strongest in *H* calculated from HbR. Together, these results show that scale-invariance in fNIRS data, particularly taken from the deoxygenated hemoglobin signal, is sensitive to differences in task difficulty and cognitive load previously identified in fMRI and EEG studies [15,22,27].

## 2. Method

This paper uses a dataset previously reported in Meidenbauer et al. (2021).

### 2.1. Participants

Seventy adults participated in this study. Participants were recruited from the Chicago area. Participants were only excluded from participating if they did not have normal or corrected-to-normal visual acuity or reported a history of neurological disorders. Participants gave written informed consent before participation and experimental procedures were approved by the University of Chicago’s Committee for Institutional Review Board (IRB). Participants were compensated $26 or 2 units of course credit, plus a performance-based bonus of up to $10. The full procedure included approximately 15 minutes of additional study elements related to a video intervention that were separate from the current work. The full study procedures lasted between 75 and 90 minutes. From the original 70 participants, 8 participants were excluded from all data analysis due to technical issues, participant non-compliance with the task procedures, or low-quality data. For this particular analysis, 10 additional participants were excluded due to insufficiently high data quality as defined by the structured noise index (SNI; see section *Exclusion of participants based on SNI*), leading to a final sample of 52 participants. Of the 52 participants analyzed here, 26 were male and 26 were female, and the mean age was 24.5 years (SD = 6.9 years).

### 2.2. fNIRS Data Acquisition

fNIRS data were acquired using a continuous-wave NIRx Nirsport2 device (NIRx Medical Technologies, LLC) and NIRx acquisition software Aurora 1.2 at a sampling rate of ∼4.4Hz. Near-infrared light of two different wavelengths (760 and 850 nm) was used to detect the concentration change of oxygenated hemoglobin (HbO) and deoxygenated hemoglobin (HbR). There were 16 sources and 16 detectors used, creating 43 channels in total, 33 channels covering the bilateral frontal cortex and 10 channels covering the right parietal cortex. Because of the low quality of parietal data collected, the following analysis focuses on data collected in the frontal region (**Fig. 1**; See section *Exclusion of noisy channels based on Structured Noise Index*). The montage was created using fOLD (fNIRS Optodes’ Location Decider;[30]), which allows placement of optodes in the international 10-10 system to maximally cover anatomical regions of interest. The AAL2 (Automated Anatomical Labeling; [31]) parcellation was used to generate the montage and provide as much coverage of the prefrontal cortex (PFC) as possible, including bilateral superior and inferior frontal gyri.

**Figure 1.**
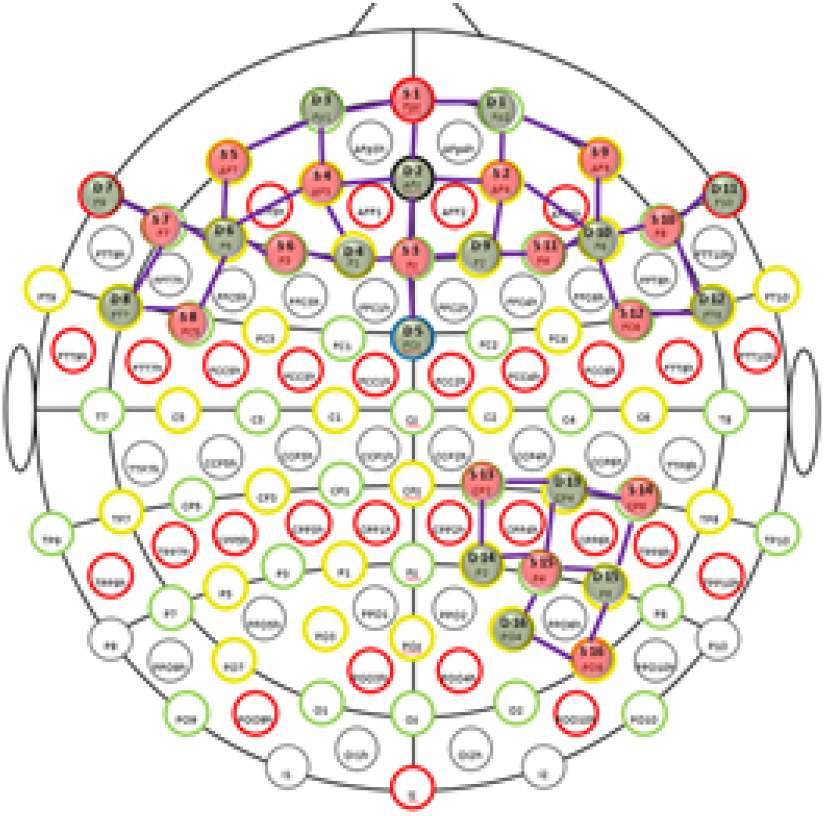
Locations of the channels in the International 10/20 coordinates. Only 33 channels covering bilateral frontal cortex are used in *H* analysis

### 2.3. N-back Task

Experimenters first took participants through step-by-step instructions of the N-back task before they began practice. Participants were told that they would see a sequence of short words that are separated by brief fixations (small circles), and that a word would be presented every 2 seconds which should be compared to the word “N” trials back. In the current study, N = 1, 2, or 3. Participants were instructed to press the “m” key every if the current word matched the word N trials back, and to press the “n” key if the current word did not match the word N trials back [**Fig. 2**]. Blocks began by first displaying the N-back level for that round and a fixation cross (5 seconds). Each task block contained a 15-length pseudorandom sequence of two words, presented for 2 seconds each (total of 30 seconds), followed by 20 seconds of rest. The length of each block was 55 seconds, and with 18 blocks the total length of the N-back task was approximately 16.5 minutes. To suppress sequence memory formation, the two words used in each block were randomly selected from the eight-word pool (‘WHAT’, ‘HOW’, ‘WHEN’, ‘WHY’, ‘WHERE’, ‘WHO’, ‘THAT’, ‘BUT’), except during the first practice, in which the same two words (“AXE” and “BOX”) were used. The sequence of two words was determined using an m-sequence (base = 2, power = 4; thus, one word appeared eight times, and the other word appeared seven times; [32–34] to suppress autocorrelation. In all cases, words were presented in white text on a black background.

**Figure 2.**
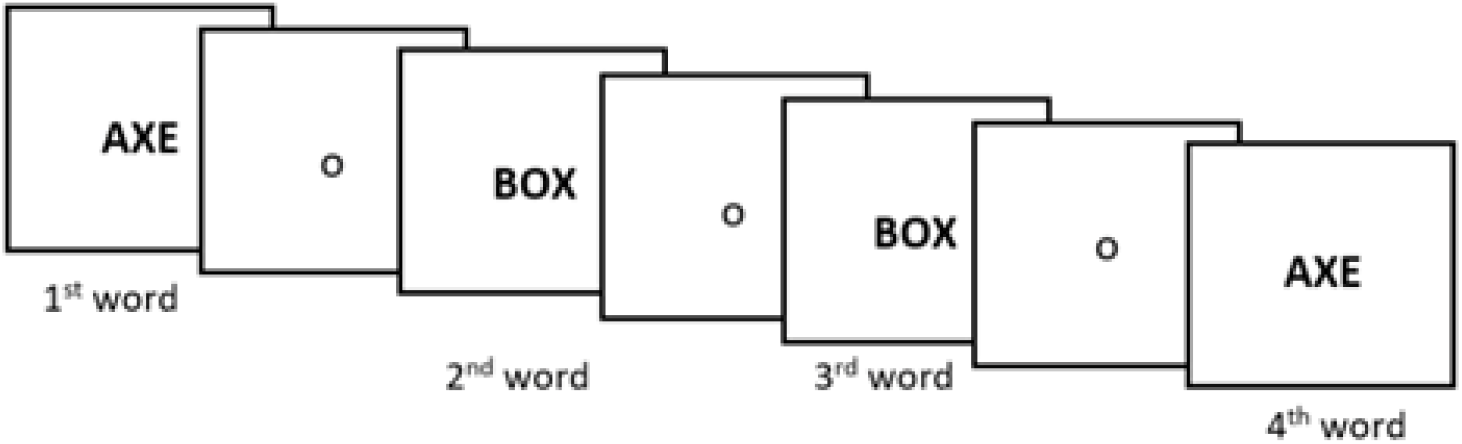
The ‘N-back’ task paradigm. Participants were required to respond whether the current word matched the word N trials back (N=1,2,3). Task difficulty and cognitive load increases with larger N.

### 2.4. Study Procedures

After providing informed consent, experimenters measured the participants’ head to determine cap size and placement, then began to set up the cap while participants were taken through task instructions and given an opportunity to practice the N-back task. The first round of N-back practice consisted of 9 blocks. In this first practice, accuracy feedback was provided on a trial-by-trial level as well as at the end of each block. Participants completed 3 blocks of 1-back, then 3 blocks of 2-back, then 3 blocks of 3-back, and then one more round of 1-back, 2-back, and 3-back. After the first round of practice, the cap was placed on the participants’ head, moving hair as needed to provide clear access to the scalp for the sources and detectors. Cap alignment was verified based on the international 10-20 location of Cz [35]. fNIRS data were calibrated and checked for quality before proceeding. If any channels were not displaying sufficiently high-quality data, placement and hair-clearing were performed again. After the fNIRS cap was set up, participants began the second round of practice designed to mimic the conditions of the real task more closely. In this practice, participants performed a single block of 1-back, then 2-back, then 3-back, without trial-by-trial feedback. The main N-back task involved 18 blocks, with 6 blocks of each N-back level, pseudorandomly presented. After completing the experiment, the cap was removed, and participants completed a demographics questionnaire. All experimental procedures were coded and presented using PsychoPy [36]. N-back experiment code can be accessed at https://osf.io/sh2bf/

Participants received a performance-based bonus during the main round (18 blocks) of N-back task. The bonus was defined as an additional 40 cents per block if performance > 90%, an additional 30 cents per block if performance > 80%, and an additional 20 cents per block if performance > 60%. Performance under 60% did not yield a cash bonus in this study. Participants were informed of their performance on each block and total bonus directly following the 30 seconds of task.

### 2.5. Data Analysis

All code used in pre-processing, *H* calculation, and statistical analysis can be accessed at: https://osf.io/kt5cx/

#### 2.5.1. fNIRS signal preprocessing

fNIRS data were first loaded into the HOMER2 software package [37] for visual inspection of overall data quality (at the level of the participant). This was done by examining the power spectral density plots for all channels to identify the presence of a cardiac oscillation (typically ∼1 Hz; [38], which indicates that the optical density signals are successfully coupled with a physiological hemodynamic response [39]. This method was used to do a first pass evaluation. For fNIRS data pre-preprocessing, this study used the Brain AnalyzIR Toolbox [40], and first converted raw light intensity values into optical density signals. Then, optical density signals were transformed into concentration changes of HbO and HbR values based on the modified Beer-Lambert law [41].

#### 2.5.2. Hurst exponent calculation on HbO and HbR data

To calculate the *H* value, the key index of interest in this study, we used the Detrended Fluctuations Analysis algorithm, which was developed for fMRI *H* analysis (DFA) [22,42]. We adopted the DFA algorithm to derive *H* from fNIRS signals in our analysis as fNIRS measures the same biological signal as fMRI.

DFA measures the power in HbO/HbR fluctuations for different time windows of the data, formulated as F(n) as a function of time in windows of length n. The Hurst exponent is equal to the slope α of a linear fit between the log-transformed F(n) and n, with α = 1 indicating a perfectly fractal signal. The length of time window, n, in our analysis was set as the full length of rest, which was 20s, with which the sampling rate of ∼4.4 Hz, yielded 87 samples/block. We kept this time window the same between task and rest so the two could be directly compared. As the task-evoked hemodynamic response occurs on a delay after the onset of the underlying neural activity, the last 20s of the 30s window of task was used in the DFA calculation.

As each N-back experiment condition had 6 blocks (18 blocks in total), including both task and rest, after the DFA calculation, each participant had 18 *H* values for task and 18 for rest (6 for each condition) for each of the 33 channels. Further visualization and statistical analysis were based on these *H* values, the key index of interest.

#### 2.5.3. Exclusion of noisy channels based on Structured Noise Index (SNI)

Before conducting statistical analysis, we first examined data quality based on the Structured Noise Index (SNI), calculated from the Brain AnalyzIR toolbox. SNI was calculated for each channel, for each participant, and is a useful tool in capturing the systematic noise across channels and participants. The SNI is defined as the ratio of the variance of the full data trace to the variance of the auto-regressively whitened trace of the same data and reflects the ratio of structured (colored) noise in the data due to various physiological processes to the uncorrelated (white) noise. This approach was inspired by the spatial SNI method described in [43] and applied to the fNIRS time signals in this work. This step was important as fNIRS signals are sensitive to superficial physiological noise (e.g., hair) and participants have varied levels of interference based on the color and texture of their hair [40]. When SNI is less than 2, it indicates that the data from this channel are very noisy and not appropriate for further analysis, and hence, we defined channels with SNI less than 2 as ‘bad’ channels.

Based on this criterion, we found that, out of total 43 channels, participants had on average 6.94 bad channels for HbO (16%) and 9.35 bad channels for HbR (22%), and that these bad channels mainly centered around parietal regions, where signals are more likely to be obstructed by hair. In this dataset, a large number of participants (46 out of 52) had one or more channels marked as “bad” in those covering the right parietal cortex (channels no. 33-43 in **Figure 1** and **Figure 3**). Therefore, in the following analysis, we excluded the 10 channels in the right parietal cortex, resulting in a total of 33 channels for analysis.

**Figure 3.**
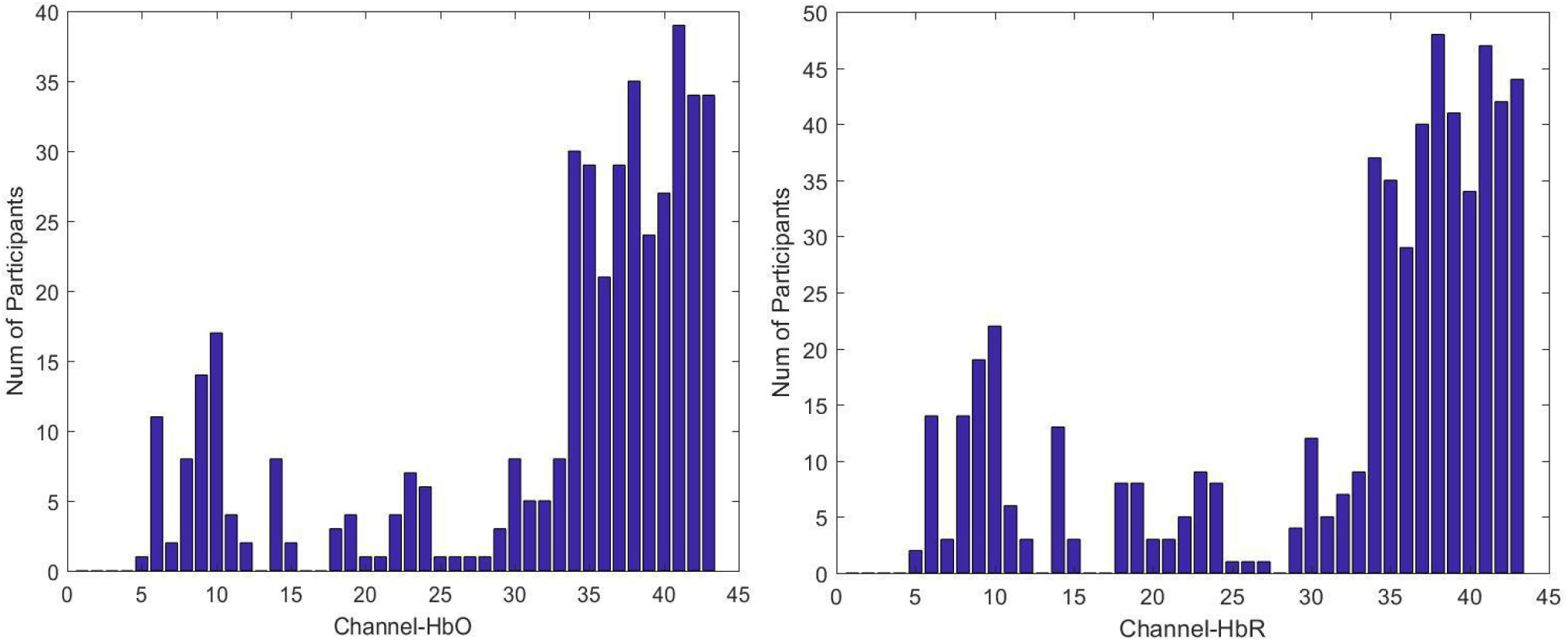
Number of Participants with each channel (1-43) marked as bad. The number of participants reporting the channel as ‘bad’ were calculated based on channel SNI < 2. Left figure for HbO; Right figure for HbR. Channels 33-43 reflect 10 channels over parietal cortex that were excluded from future analysis.

#### 2.5.4. Exclusion of participants based on SNI

Additionally, after excluding those 10 channels in the parietal cortex, 10 participants’ data were found to have many poor-quality channels (shown in **Figure 4**). This was defined by participants whose count of bad channels exceeded 1 SD (3.0 bad channels for HbO; 3.4 bad channels for HbR) from the average number of bad channels (2 bad channels for HbO; 3 bad channels for HbR). Following this criterion, participants who either had 5 bad channels for HbO or 6 bad channels for HbR were excluded from further analysis. On average, these 10 outlier participants had 8 (24%) bad channels for HbO and 9 (27%) bad channels for HbR.

**Figure 4.**
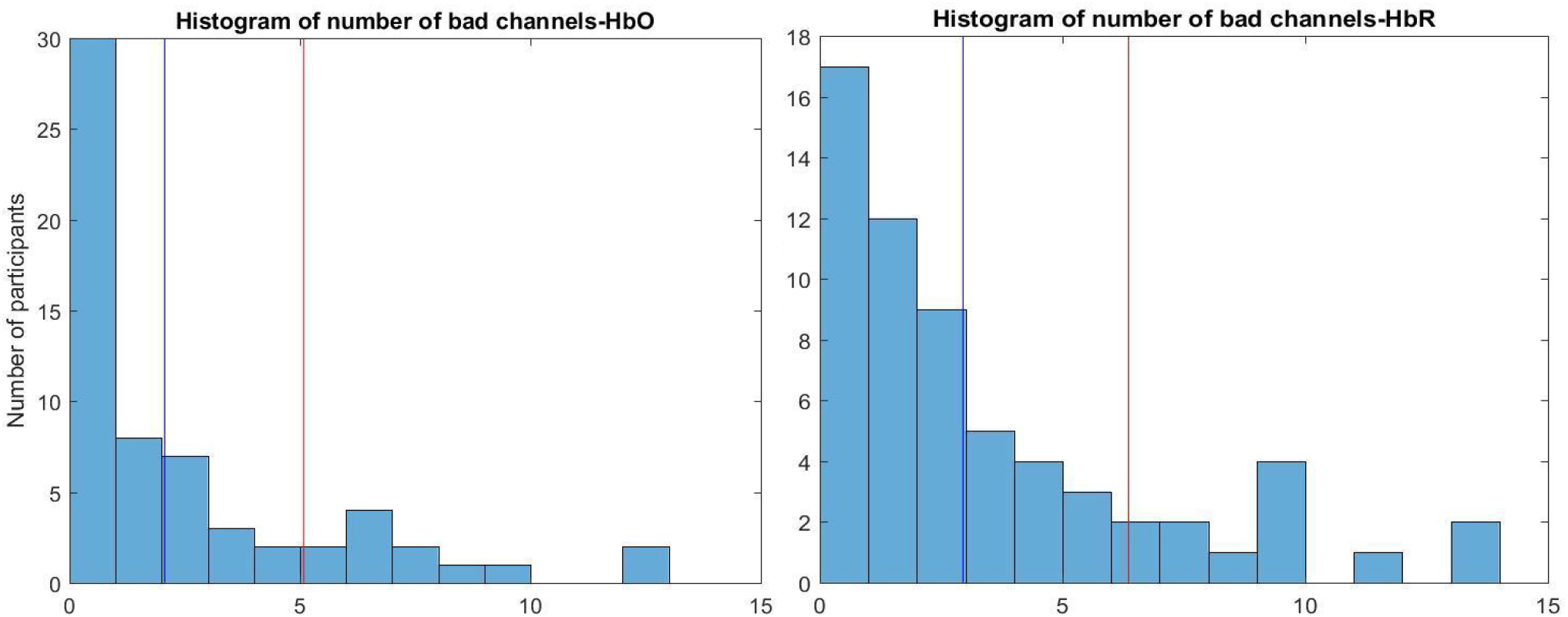
Number of ‘bad’ channels per participant after excluding parietal channels. Histograms show the number of ‘bad’ (SNI <2) channels (out of 33) each participant had after removing the 10 parietal channels for HbO (left) and HbR (right). The blue line represents the mean number of bad channels across participants and the red lines indicate 1 SD from the mean.

As an additional check, after calculating the Hurst exponent, we plotted average *H* across all task sessions by channels for each participant. As *H* is a global signal across the whole brain, we expected moderate to high levels of similarity across channels [22,44]. Large fluctuations between channels of the same participant in the same run suggest poor data quality, potentially stemming from hair thickness variations. Shown in **Fig. 5**, we could see that compared to ‘good’ participants (lower panel B), ‘bad’ participants (upper panel A) generally had more variable responses across channels that were very irregular and unexpected, which further justified this SNI based participants exclusion step.

**Figure 5.**
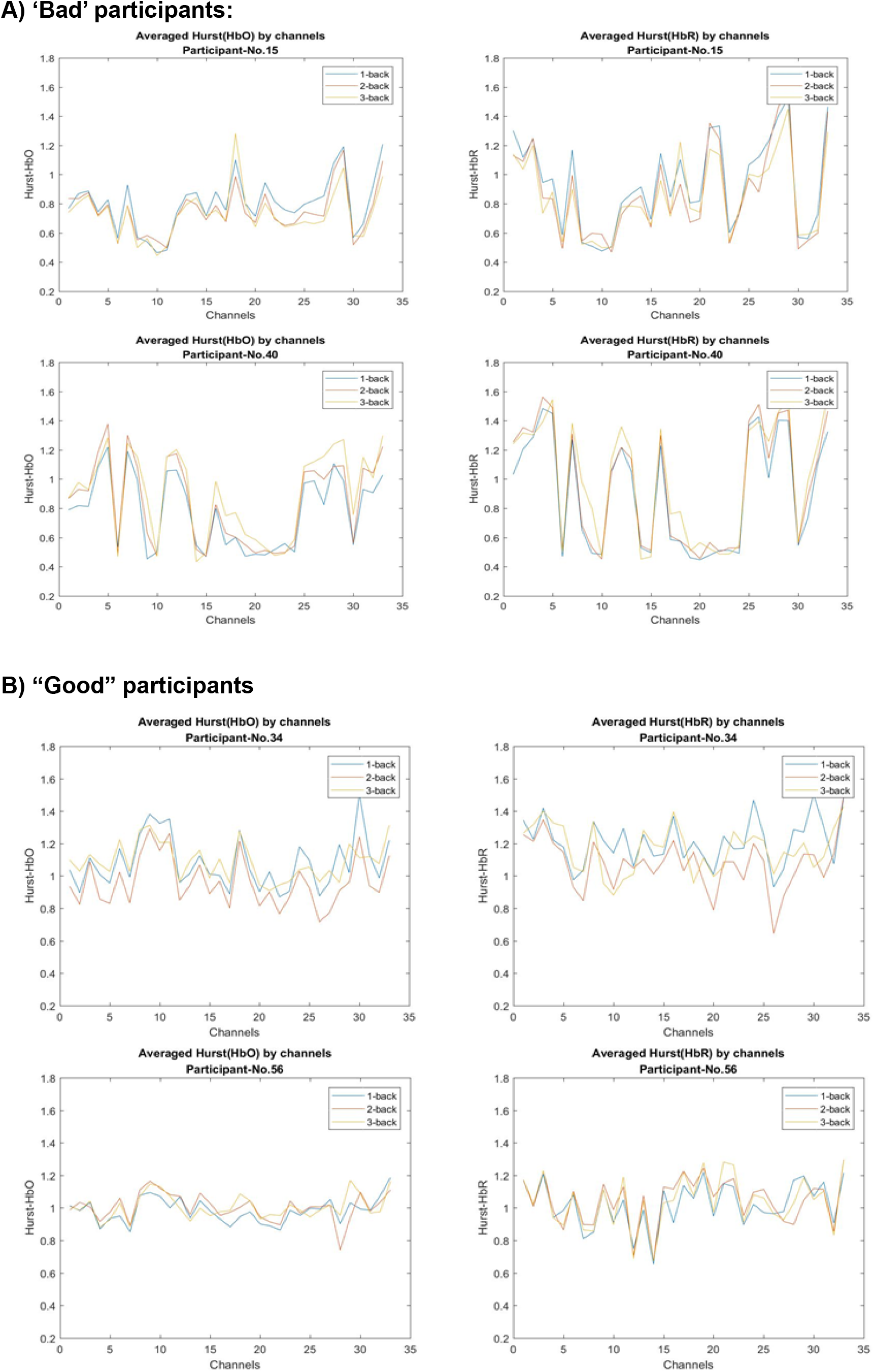
Averaged *H* value by channels for participants with good and bad data quality. *H* score of each 20 segments was averaged for each block type for each participant. The upper 4 figures (panel A) showed participants marked as ‘bad’ (left for HbO, right for HbR); the lower 4 (panel B) showed participants which were not marked as ‘bad’ (left for HbO, right for HbR).

#### 2.5.5. Regressing SNI from H values

We further regressed the SNI out from the *H* value of each channel, which accounts for different degrees of noise across channels, to remove the systematic interference of noisy data. The value of *H* after SNI being regressed out was used in further analysis.

#### 2.5.6. Accounting for Motion-related Artifacts

Lastly, as the scale-invariance of a data time series can be influenced by the presence of large fluctuations not due to the underlying biological signal, such as motion-related artifacts [45], we evaluated the presence of such artifacts across conditions and for task and rest separately to ensure these were not driving any results. For example, if participants in the study systematically demonstrated more head movement more during rest blocks than during task blocks, this would lessen the interpretability of our *H* results. To do this, the raw data were first segmented into the same 20 second blocks used in the DFA calculation and statistical analysis. Next, statistical outliers in the 20 seconds of data were identified by calculating the innovations model for each segment using the function nirs.math.innovations() in the Brain AnalyzIR Toolbox [40]. The innovations in each time series reflect the uncorrelated (whitened) signal after filtering using the autoregressive model. Here, each 20 second time series for the 33 channels used in the Hurst analyses for each participant was fed into the function, and the maximum model order was set to 20. This was chosen based on the recommendation that the maximum model order is at least 4 times the sample rate (here ∼4.4 Hz).

The output of this function is an innovations time series for each channel for each segment. Subsequently, the number of studentized outliers were calculated (outliers defined as *p* < 0.05). The counts of these outliers across all 33 channels for each segment and each participant were then saved. The average number of statistical outliers by segment type (all task, all rest, 1-back, 2-back, and 3-back) across are presented in **Table 1**. Paired t-tests (mimicking the statistical tests on averaged *H* values, detailed below) were also conducted to compare the number of outliers/artifacts by condition. The results of these tests are also presented in Table 1. The number of outliers was generally quite low: out of ∼2870 data points reflecting time (∼87 samples) x 33 channels, the average number of outliers participants had for each condition was between 182 and 200, or roughly 6.3% to 6.9% of the data. The counts were also very similar across conditions [**Table 1**]. For those comparisons which did show a significant difference between conditions, the relative number of outliers were in the opposite direction of what would be expected if these motion artifacts were affecting the *H* results. This suggests that these outliers (which exceed what is a reasonable change to be expected from the biological signal of interest) are not responsible for differences in *H* values across conditions.

**Table 1:** Counts & comparisons of average number of motion related artifacts. The average number of motion artifacts were calculated for each participant in each 20 second block type (all task, all rest, 1-back, 2-back, and 3-back). Each average count is out of 2871 potential data points (33 channels x 87 samples). *M* and *SD* calculated over the 52 participants’ average motion artifacts in each block type. *T*-tests reflect paired (within-subjects) comparisons of motion artifact counts in each block type.

| **Counts** | **HbO time series**<br>**M (SD)** | **HbR time series**<br>**M (SD)** |
| --- | --- | --- |
| **Task** | 183.6 (71.8) | 197.5 (65.0) |
| **Rest** | 184.1 (71.6) | 198.8 (64.1) |
| **1-back Task** | 182.7 (71.8) | 200.1 (67.3) |
| **2-back Task** | 183.1 (72.1) | 196.1 (64.4) |
| **3-back Task** | 185.5 (71.8) | 196.2 (63.3) |

| ***Pairwise T-tests*** | **HbO** | **HbR** |
| --- | --- | --- |
| **Task vs. Rest** | $t(51) = -0.48, p = 0.63$ | $t(51) = -1.78, p = 0.08$ |
| **1-back vs. 2-back** | $t(51) = -0.36, p = 0.72$ | $t(51) = 2.57, p = 0.01$ |
| **2-back vs. 3-back** | $t(51) = -1.41, p = 0.16$ | $t(51) = -0.07, p = 0.94$ |
| **1-back vs. 3-back** | $t(51) = -1.77, p = 0.08$ | $t(51) = 2.11, p = 0.04$ |

#### 2.5.7. Statistical Analysis on Average *H*

Using the cleaned averaged *H* values with SNI regressed out, planned pairwise *t*-tests between task and rest were conducted on HbO and HbR separately. Next, two repeated measurements ANOVAs (separately for HbO and HbR) were conducted on average *H* value with N-back condition (1-back, 2-back, or 3-back) as a within-subjects factor. Lastly, we also examined planned pairwise *t*-tests comparing each of the three N-back conditions. These analyses were carried out using R version 3.5.1 [46].

#### 2.5.8. PLS Analysis

In addition to statistical analyses on the Hurst values averaged across channels, partial least squares (PLS; [47–49]; https://www.rotman-baycrest.on.ca) analyses were also conducted using Matlab v 2018b. PLS is a data-driven, multivariate statistical technique which identifies the relationship between two sets of variables. In neuroimaging research, PLS is often used to find the relationship between neural activity at different spatial locations (e.g., voxels, or ROIs in fMRI data, electrodes in EEG data, channels in fNIRS data) and the task design (e.g., experimental conditions or grouping variables). In the current work, a Task PLS was conducted to examine *H* by N-back level across channels.

The partial least squares analysis relies on the singular value decomposition (SVD) of a covariance matrix. In the Task PLS analysis, the input for SVD is a matrix of the *H* values for each channel by condition (N-back level) that are averaged across participants (i.e., matrix of 33 channels x 3 N-back levels). Running an SVD on this 33 × 3 covariance matrix (R) decomposes it into three matrices: R = UΔV^T^, where the 3×3 matrix U represents the decomposition of R in N-back condition space, the 33 × 3 matrix V represents the decomposition of R in neural activity optode space, and Δ is the 3×3 diagonal matrix of singular values that quantifies the weighting of each of the singular vectors (i.e., columns in U and V’). These linear decompositions which maximize the covariance between brain activity (*H* values) and task design (N-back level) are referred to as latent variables (LVs). These LVs are calculated in order of magnitude of cross-block covariance explained and are mutually orthogonal, so the first latent variable (LV 1) explains the greatest proportion of cross-block covariance, the second latent variable (LV 2) explains the second most proportion, etc.

Ten thousand permutation tests were performed to obtain *p*-values for each latent variable and 10,000 bootstrapped samples with replacement were created to generate the 95% confidence intervals for variable loadings. The bootstrap ratios (calculated as salience[weights] / standard error[reliability]) measure the reliability of the relationship at each channel, and a larger bootstrap ratio indicates a stronger and/or more consistent contribution to the LV. In this study, channels with bootstrap ratios larger than +2 or smaller than −2 were determined to be statistically significant as these bootstrap ratios can be interpreted as z-scores.

## 3. Results

### 3.1. Statistical Analysis on Averaged Hurst Exponents

For the task and rest comparison, planned pairwise *t-tests* showed that compared with the rest condition, averaged *H* scores were significantly lower in the task conditions for both HbO (Task M = 0.83, SD = 0.15, Rest M = 0.96, SD = 0.15, *t*(51)=11.76 *p*<0.001) and HbR (Task M = 0.88, SD = 0.17, Rest M = 0.94, SD = 0.16; *t*(51)=7.72, *p*<0.001). These results are shown in **Figure 6**.

**Figure 6.**
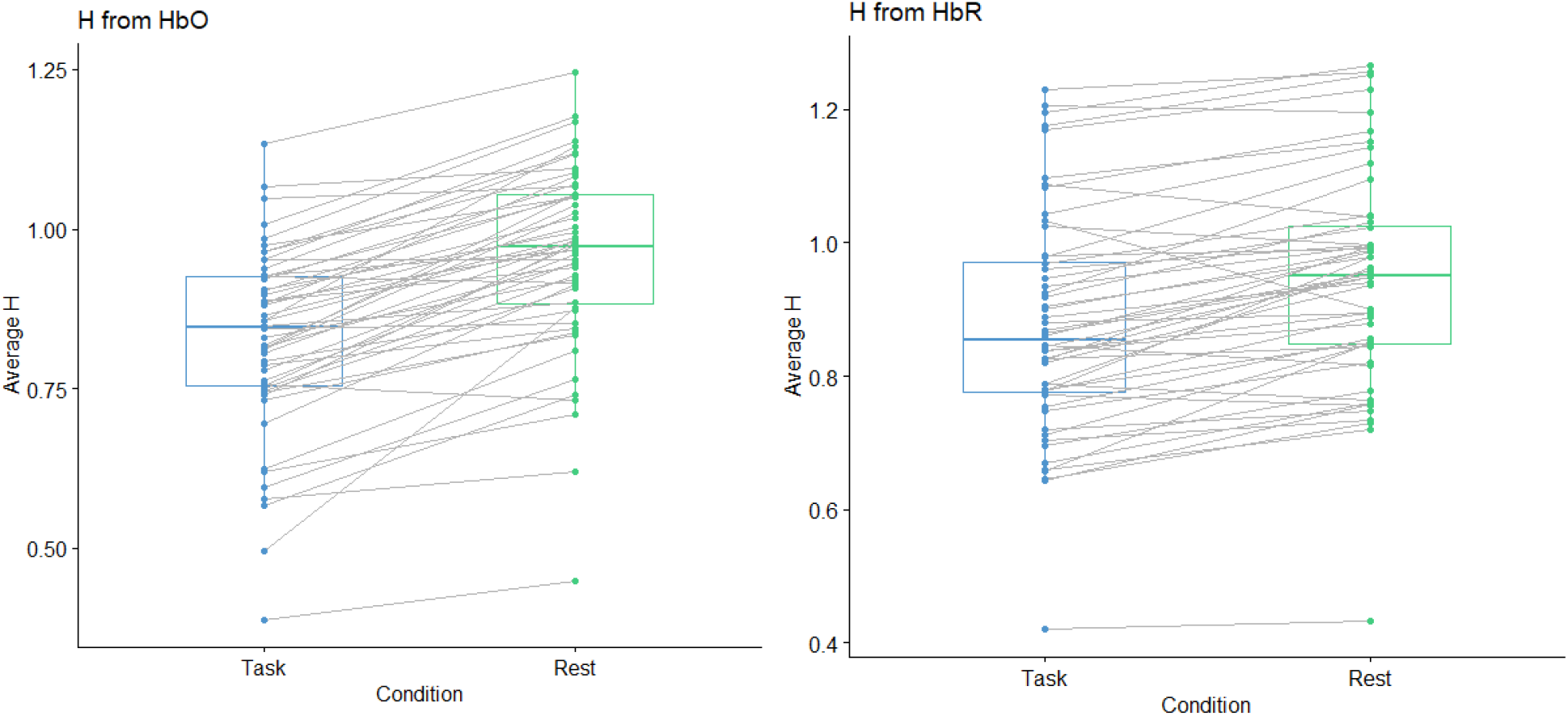
Boxplot showing within-subjects effects for Averaged *H* for Task vs. Rest. Gray lines connect *H* values for rest and task for each participant.

For the N-back conditions comparison, where N-back condition is a within-subjects factor, repeated measures ANOVAs on averaged *H* by both HbO and HbR showed a significant omnibus ANOVA for *H* extracted from HbR (*F*(2,102)=3.78, *p*=0.026), but not from HbO (*F*(2,102)= 1.92, *p*=0.153). In both *H* extracted from HbO and HbR, 2-back had the lowest value (HbO M = 0.82, SD = 0.15; HbR M = 0.82, SD = 0.16) of *H*, followed by 3-back (HbO M = 0.83, SD = 0.15; HbR M = 0.88, SD = 0.16) and then 1-back (HbO M = 0.82, SD = 0.15; HbR M = 0.82, SD = 0.16.

For planned pairwise *t*-tests, *H* extracted from HbR was significantly different between the 1-back and 2-back conditions (*t*(51) = 2.62, *p* = 0.011), and marginal for *H* by HbO (*t*(51) = −1.80, *p* = 0.078). Pairwise *t*-tests between 2-back and 3-back were marginal for *H* extracted from HbR (*t*(51) = −1.80, *p* = 0.078). No other *t*-tests were statistically significant for either *H* extracted from HbO or HbR (all p > 0.24).

### 3.2. PLS Results

Task PLS analyses looking at channel-level *H* by N-back condition were run separately on *H* extracted from HbO and from HbR time series. The first latent variable (LV 1) from the analysis with *H* from deoxyhemoglobin concentrations (HbR) was significant and explained 77% of the crossblock covariance (*p* = 0.005). LVs 2 and 3 in this analysis were not significant (all ps > 0.4). For the significant LV 1 in *H* from HbR, 11 mostly medial-frontal channels (#1, #5, #6, #7, #8, #11, #12, #13, #14, #20, and #25) showed stable changes in scale-invariance by N-back level, indicated by bootstrap ratios with absolute values greater than 2 [Table 2; Figure 7]. LV 1 from the analysis of *H* calculated from oxyhemoglobin concentrations (HbO) was marginal (*p* = 0.066) and also explained 77% of the crossblock covariance. LVs 2 and 3 in this analysis were not significant (all *ps* > 0.17). For LV1, 2 channels in the medial superior frontal gyrus (#1 and #2) and 2 in the left inferior frontal gyrus (#22 and #23) showed differences in *H* by N-back level. In all cases, the significant bootstrap ratios were greater than 2 (and none smaller than −2), indicating the relationship between N-back level and *H* was in the same direction. In *H* calculated from HbO and HbR, the first latent variable from each was driven primarily by the contrast between 1-back and 2-back, with higher *H* found during the 1-back task relative to the 2-back task. [Figure 7]

**Table 2.**
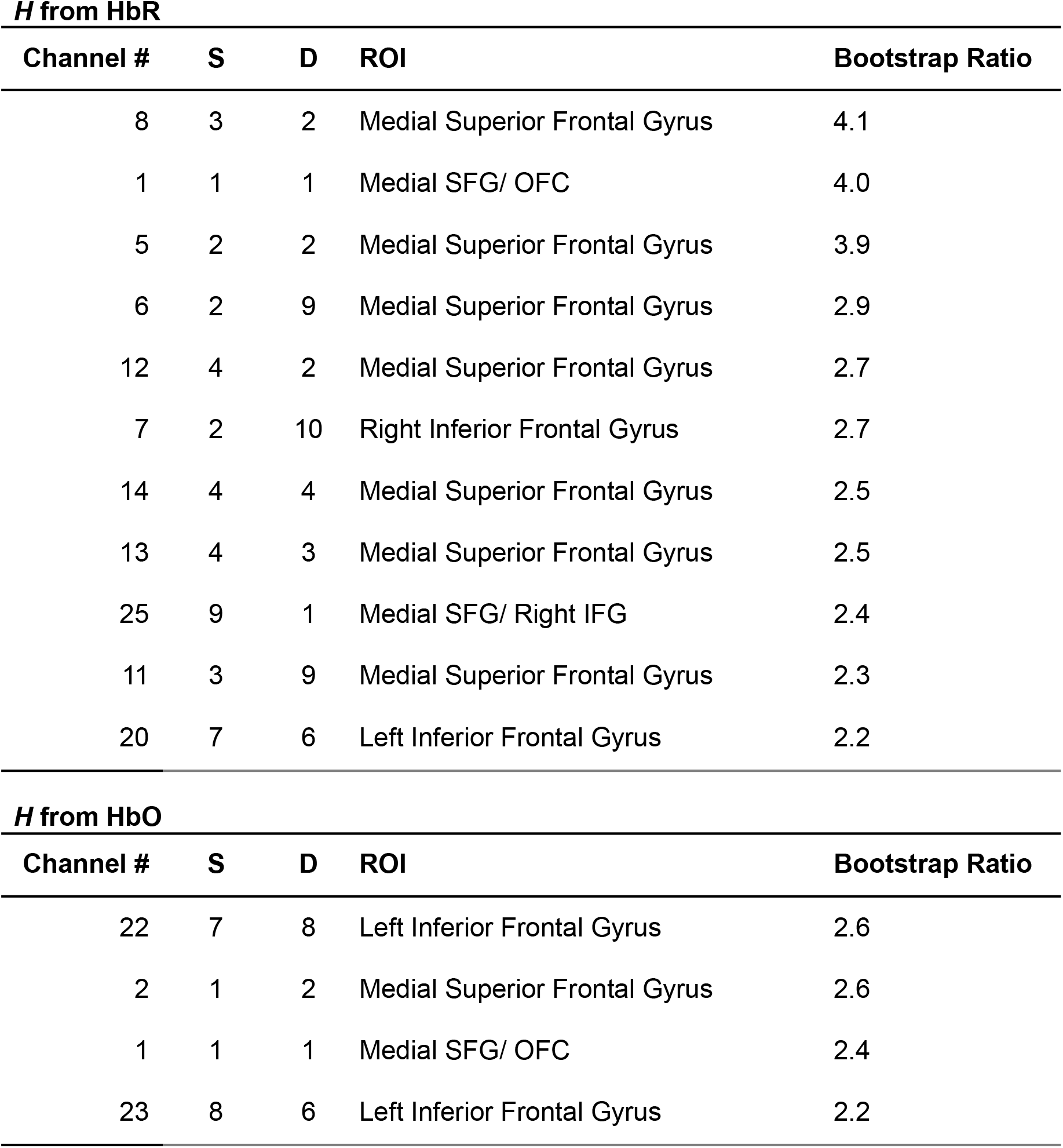
Significant Channels for Task PLS LV 1. Top: Results for *H* from deoxyhemoglobin (HbR), Bottom: Results for *H* from oxyhemoglobin (HbO). Channel number based on source (S) - detector (D) pair. ROI label defined by maximal coverage of talairach daemon ROI. Channels ordered by size of bootstrap ratio. Bootstrap ratios > |2| were considered significant.

| Channel # | S | D | ROI | Bootstrap Ratio |
| --- | --- | --- | --- | --- |
| 8 | 3 | 2 | Medial Superior Frontal Gyrus | 4.1 |
| 1 | 1 | 1 | Medial SFG/ OFC | 4.0 |
| 5 | 2 | 2 | Medial Superior Frontal Gyrus | 3.9 |
| 6 | 2 | 9 | Medial Superior Frontal Gyrus | 2.9 |
| 12 | 4 | 2 | Medial Superior Frontal Gyrus | 2.7 |
| 7 | 2 | 10 | Right Inferior Frontal Gyrus | 2.7 |
| 14 | 4 | 4 | Medial Superior Frontal Gyrus | 2.5 |
| 13 | 4 | 3 | Medial Superior Frontal Gyrus | 2.5 |
| 25 | 9 | 1 | Medial SFG/ Right IFG | 2.4 |
| 11 | 3 | 9 | Medial Superior Frontal Gyrus | 2.3 |
| 20 | 7 | 6 | Left Inferior Frontal Gyrus | 2.2 |

***H* from HbO**
| Channel # | S | D | ROI | Bootstrap Ratio |
| --- | --- | --- | --- | --- |
| 22 | 7 | 8 | Left Inferior Frontal Gyrus | 2.6 |
| 2 | 1 | 2 | Medial Superior Frontal Gyrus | 2.6 |
| 1 | 1 | 1 | Medial SFG/ OFC | 2.4 |
| 23 | 8 | 6 | Left Inferior Frontal Gyrus | 2.2 |

**Figure 7.**
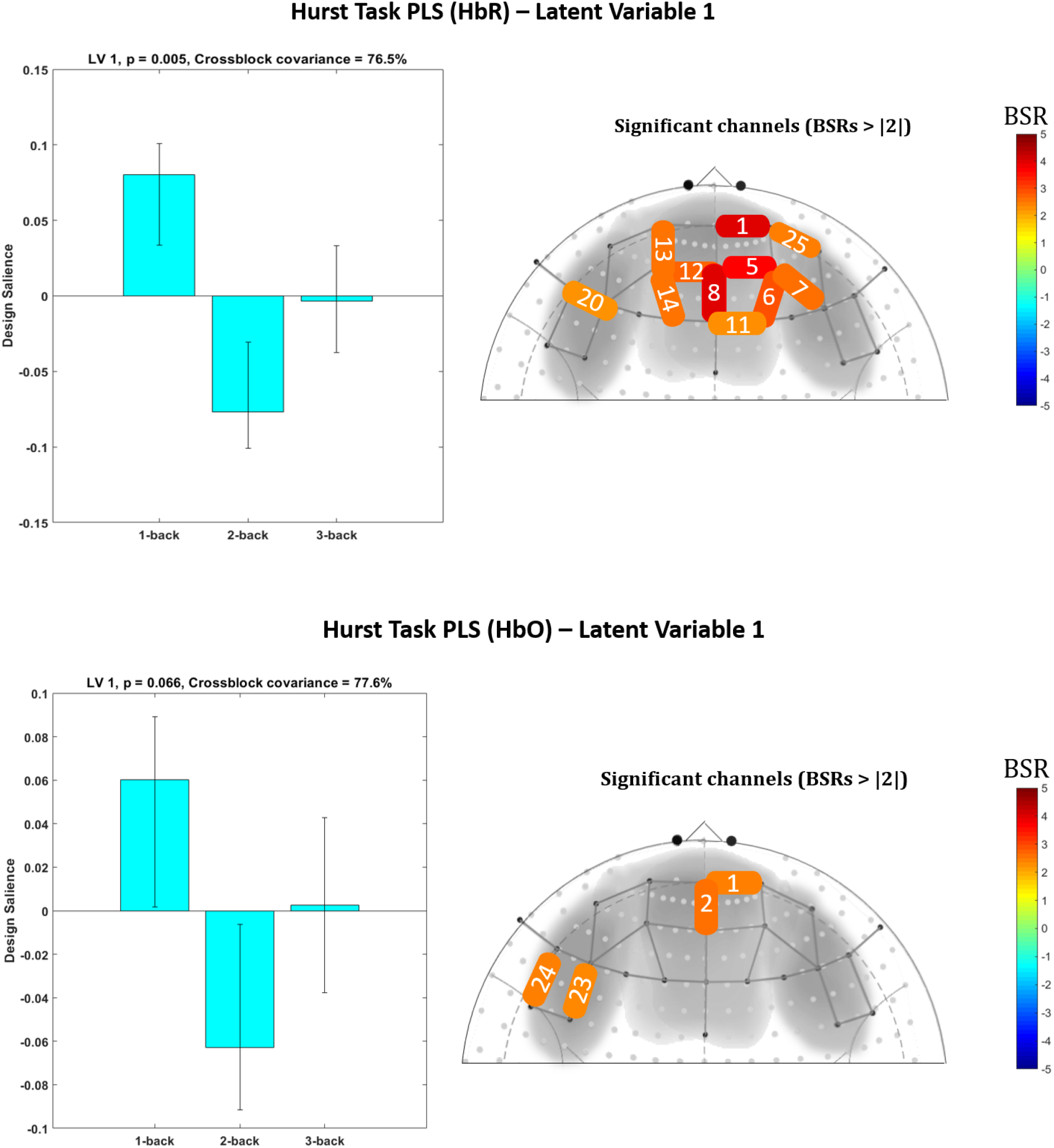
LV 1 demonstrated an N-back load-dependent relationship with *H*. extracted from deoxyhemoglobin (HbR; Top Panel) and oxyhemoglobin (HbO; Bottom Panel) concentrations. The left side plots show the relation between *H* and N-back level. Error bars are 95% confidence intervals around the mean design salience value. The right panel shows channels (labeled by number), which had bootstrap ratios (BSR) > |2|.

## 4. Discussion

Previous research suggests that when people are performing a task of high cognitive demands, compared with rest, the temporal property of their brain signals will be less scale invariant as quantified by a lower *H* score. In addition, this suppression of scale invariance is found across the whole brain [15,28,50] and is unidirectional [22]. However, whether this signature could be extracted from fNIRS data was unclear. The present study is the first to apply a scale-invariance (Hurst exponent, *H*) analysis to measuring cognitive load with fNIRS. Consistent with previous neuroimaging research, we found that task and rest conditions significantly differed by their average *H* calculated from both oxyhemoglobin (HbO) and deoxyhemoglobin (HbR) concentrations changes of the fNIRS signal. Compared with rest, average *H* for task was significantly lower, which suggests a higher level of cognitive effort and difficulty while performing the N-back task relative to rest. For N-back condition, a more subtle manipulation of cognitive load, pairwise *t*-tests also showed a significant difference between 1-back and 2-back by *H* calculated from HbR. This was marginally significant with *H* from HbO, where again, *H* during 2-back had lower values than 1-back. While these effects with HbO were not statistically significant, they do demonstrate convergence with the pattern observed in HbR and in comparing task vs. rest.

The N-back level results from averaging across all 33 channels were further supported by those yielded from the partial least squares analysis. Specifically, in the Task PLS, which examines differences in *H* across channels and N-back level, the first latent variable from HbR demonstrated a robust effect of higher *H* in 1-back vs. 2-back across 11 frontal channels. Though the effect was weaker in *H* calculated from HbO, the same pattern was found in this analysis’ LV 1. These results provide complementary evidence for an effect of cognitive load and task difficulty on *H* derived from fNIRS. Additionally, they suggest that while *H* is believed to be a relatively global brain signature [22], greater sensitivity can be achieved by adopting a multivariate technique which takes the full data (including channels) as input.

It is worth noting that both in the pairwise comparisons of average *H* by N-back condition and the Task PLS analysis, this study only yielded significant results for the 1-back vs. 2-back comparison. In contrast, the 2-back vs. 3-back comparison was marginally significant for *H* by HbR (not with HbO), and all the other *t*-test results for 3-back condition (3-back vs. 1-back and 3-back vs. 2-back) were not significant. However, as observed in Meidenbauer et al. (2021) which used this same dataset, performance in 3-back condition was generally low and was highly variable.

The non-significant results for the 3-back condition have also been shown in previous studies, indicated by a non-linear effect of N-back load [51–53]]. Researchers have argued that if a task is too difficult, people may disengage from it or simply “give up”, since it exceeds one’s capability [52,54]. Following this argument and based on the results reported in Meidenbauer et al. (2021), we infer that in this study, the 3-back condition might not be reflecting the highest cognitive load across all participants. Results of a multivariate analysis including performance and standard task-evoked activation in this same dataset further support this idea [29] Thus, the large individual differences and generally poor performance of the 3-back condition may explain the non-significant results in this study with *H* and should be examined in more detail in future work.

### Data preprocessing pipeline for H analysis with fNIRS

As the first study exploring the application of scale invariance in fNIRS, this study developed a basic pipeline for data preprocessing and *H* analysis with fNIRS. (All analysis scripts can be accessed at: https://osf.io/kt5cx/) To check the quality of our data, we used the SNI (Structured Noise Index) to measure the systematic noise across channels. We subsequently excluded participants who had a high number of low SNI channels and regressed out SNI from the Hurst exponent. Both steps were supported by visualizations as effective in detecting poor data quality, finding ‘outliers’, and removing systematic noise. These steps were adopted for two primary reasons. First, superficial physiological noise is structured in fNIRS and can differ by person according to how their hair may obstruct signals [40,55]. Second, scale invariance is generally a whole brain index that, when examined across channels, could be biased by channel-level differences in superficial noise. To further rule out the influence of motion-related artifacts, we used an innovations model to calculate the statistical outliers of the time series (i.e., the signal variations which are larger than what is expected due to underlying physiological changes) and found that the motion-related artifacts did not explain *H* effects.

### Implications of the current study

This study sheds light on the reliability of scale invariance across neuroimaging modalities and on the promising future of adopting fNIRS in examining cognitive load in real life scenarios. Relative to EEG and fMRI, fNIRS is more flexible, less affected by environmental noise, and more robust to motion artifacts. fNIRS is already used to study cognitive processes in more ecological valid scenarios [24,56]. The results of this study demonstrate the effectiveness of *H* in measuring cognitive load with fNIRS and further strengthen its capability in real-world settings, such as monitoring cognitive load during driving, social interactions, or even to examine cognitive restoration during real interaction with natural versus urban environments [57].

### Limitations and Future directions

As this study is the first to demonstrate the effectiveness of *H* with fNIRS in measuring cognitive load, there are several limitations which require further investigation. First, applying scale invariance analysis in various datasets and experiment settings (especially in real-world scenarios) would be necessary in the future to further validate the effectiveness and robustness of this measure of cognitive load with fNIRS. Secondly, since this analysis measures scale invariance on a temporal scale, the time window of analysis might impact the results. In this study, we adopted 20s as the time window to match the full length of the rest session. As fNIRS has a relatively high sampling rate, 20 seconds x 4.5 Hz provided a sufficiently high number of samples for the DFA to be reliable. However, future work should examine whether and how different time window lengths might impact its effectiveness.

Due to the high correspondence between fNIRS and fMRI, we adopted the DFA algorithm from fMRI Hurst exponent analysis [22] and showed its effectiveness with the current fNIRS pre-processing pipeline. However, future work could explore other non-stationary algorithms to calculate *H* and also time windows of different lengths, which might impact the results of *H* analysis with fNIRS.

Lastly, we found that *H* calculated from deoxyhemoglobin (HbR) showed more reliable effects than did *H* from oxyhemoglobin (HbO). This may be due to the fact that HbR is more tightly coupled with the BOLD response in fMRI [1]. However, as this is the first study to look at *H* in fNIRS, it is unclear whether this is a reliable pattern or is related to the current task design, and future investigations with different tasks, contexts, and time windows would be needed to shed light on these possibilities.

## Conclusion

This study was the first to examine the Hurst exponent as an effective measure of cognitive load with fNIRS and demonstrated a basic and robust pipeline for scale-invariance analysis in fNIRS, which lays the foundation for future theoretical and practical work on cognitive load with *H* and fNIRS. Future work could further test its theoretical validity and explore its implications with fNIRS in the real world.

## Data and code availability statement

Raw data, processed data, *H* calculation code, and all statistical analysis code can be accessed at the OSF project page: https://osf.io/kt5cx/

PsychoPy N-back Experiment code can be accessed at the OSF page associated with Meidenbauer et al., *NeuroImage* (2021): https://osf.io/sh2bf/

## Acknowledgments

We thank Jaime Young, Tanvi Laktahkia, Olivia Paraschos, and Anabella Pinton for their assistance in data collection.

## Author CRediT Statement

**Conceptualization:** Kimberly L. Meidenbauer, Kyoung W. Choe, and Marc G. Berman.

**Data curation:** Chu Zhuang and Kimberly L. Meidenbauer.

**Formal analysis:** Chu Zhuang, Kimberly L. Meidenbauer, Omid Kardan, Andrew J. Stier, Kyoung W. Choe, Theodore J. Huppert, and Marc G. Berman.

**Funding acquisition:** Kyoung W. Choe and Marc G. Berman.

**Investigation:** Kimberly L. Meidenbauer and Kyoung W. Choe.

**Methodology:** Kyoung W. Choe and Theodore J. Huppert.

**Project administration:** Kimberly L. Meidenbauer, Kyoung W. Choe, and Marc G. Berman.

**Resources:** Omid Kardan, Andrew J. Stier, Kyoung W. Choe, Carlos Cardenas-Iniguez, and Theodore J. Huppert.

**Software:** Chu Zhuang, Kimberly L. Meidenbauer, Kyoung W. Choe, Carlos Cardenas-Iniguez, and Theodore J. Huppert.

**Supervision:** Marc G. Berman.

**Visualization:** Chu Zhuang and Kimberly L. Meidenbauer.

**Writing - original draft:** Chu Zhuang and Kimberly L. Meidenbauer.

**Writing - review & editing:** Chu Zhuang, Kimberly L. Meidenbauer, Omid Kardan, Andrew J. Stier, Kyoung W. Choe, Theodore J. Huppert, and Marc G. Berman.

